# Transcranial Magnetic Stimulation-induced motor cortex activity influences visual awareness judgments

**DOI:** 10.1101/848457

**Authors:** Justyna Hobot, Marcin Koculak, Borysław Paulewicz, Kristian Sandberg, Michał Wierzchoń

**Author notes:** **Correspondence:** Justyna Hobot.

## Abstract

The influence of non-visual information on visual awareness judgments has recently gained substantial interest. Using single-pulse Transcranial Magnetic Stimulation (TMS), we investigate the potential contribution of evidence from the motor system to judgment of visual awareness. We hypothesized that TMS-induced activity in the primary motor cortex (M1) would increase reported visual awareness as compared to the control condition. Additionally, we investigated whether TMS-induced motor-evoked potential could measure accumulated evidence for stimulus perception. Following stimulus presentation and TMS, participants first rated their visual awareness verbally using the Perceptual Awareness Scale, after which they responded manually to a Gabor orientation identification task. Delivering TMS to M1 resulted in higher average awareness ratings as compared to the control condition, in both correct and incorrect identification task response trials, when the hand with which participants responded was contralateral to the stimulated hemisphere (TMS-response-congruent trials). This effect was accompanied by longer Perceptual Awareness Scale response times, irrespective of the congruence between TMS and identification response. Moreover, longer identification response times were observed in TMS-response-congruent trials in the M1 condition as compared to the control condition. Additionally, the amplitudes of motor-evoked potentials were related to the awareness ratings when response congruence was taken into account. We argue that motor-evoked potential can serve as an indirect measure of evidence accumulated for stimulus perception and that longer Perceptual Awareness Scale response times and higher amplitudes of motor-evoked potentials in the M1 condition reflect integration of additional evidence with visual awareness judgment. In conclusion, we advocate that motor activity influences perceptual awareness judgments.

## 1. Introduction

Uncovering the neural processes that shape conscious content is considered a central problem in consciousness science (Faivre et al., 2017). Access to conscious content is based on the accumulation of stimulus-based evidence, prior knowledge, and biases (Dehaene, 2008; Dehaene & Changeux, 2011; Lau, 2008). We consider conscious access to be a non-dichotomous phenomenon (Fazekas & Overgaard, 2016, 2018a; Jonkisz et al. 2017; Kouider et al., 2010; Lyyra, 2019, for alternative explanations see: Del Cul et al., 2007; Sergent & Dehaene, 2004) that is reflected in awareness judgments (Anzulewicz et al., 2019). Therefore, we operationalize conscious access with perceptual awareness ratings. This approach is consistent with several major views on consciousness, including the hierarchical view (Lau & Rosenthal, 2011), the Partial Awareness Hypothesis (Kouider et al., 2010), and some current understandings of conscious access, e.g. the Multi-Factor Account of Degrees of Awareness (Fazekas & Overgaard, 2016, 2018a). Perceptual awareness judgments – like decision confidence judgments – are a type of metacognitive judgment (Lau & Rosenthal, 2011; Fleming, 2020) and can be measured on multiple scales, such as continuous visual analogue scales (Hayes & Patterson, 1921; Sergent & Dehaene, 2004) and the Perceptual Awareness Scale (PAS; Ramsøy & Overgaard, 2004). The latter requires participants to rate stimulus awareness with ratings ranging between “no experience” and “a clear experience”. PAS is considered a sensitive and exhaustive measure of stimulus awareness (Sandberg et al., 2010) and is widely used in consciousness research (Sandberg & Overgaard, 2015).

Multiple theories frame conscious access (more or less explicitly) in the context of stimulus evidence accumulation (Block, 2011; Dehaene et al., 2003; Lamme, 2010; Mudrik et al., 2016). This has bound research to experimental paradigms that manipulate stimuli characteristics; however, the physical qualities of stimuli do not fully explain the qualities of conscious access, which implies the presence of additional sources of evidence (Anzulewicz & Wierzchoń, 2018; Tagliabue et al., 2019). Several such sources have been proposed, e.g. prior expectations (Snyder et al., 2015), previous responses (Rahnev et al., 2015), or attentional engagement (Fazekas & Overgaard, 2018b). Nevertheless, these sources are associated with the early stages of awareness-related processing (e.g. Dehaene et al., 2014). Here, we present an investigation of motor-related information influence that occurs at the later stages of awareness-related processing on stimulus awareness judgment.

Out of many possible contributors, the motor system seems especially related to perception. Numerous studies have explored the action-perception loop and have shown that in tasks requiring coordination of perceptual information and action, both systems influence each other and enhance task performance (Donnarumma et al., 2017; Hecht et al., 2001). Similar conclusions have come from experiments in which coupling between perception and action was more superficial than in action-perception loop procedures (e.g. linking certain stimuli to particular response keys: Siedlecka et al., 2019; Siedlecka, Paulewicz, Koculak, 2020a). A recent study showed that visual awareness judgments are sensitive to accuracy feedback in a stimulus identification task (Siedlecka et al., 2020b). Participants reported lower awareness after an incorrect response in the previous trial, and the effect was strengthened by trial-by-trial accuracy feedback.

Nevertheless, a couple of studies have presented a more immediate effect that shapes the experience of just-presented stimuli. Several studies have shown a consistent effect of identification tasks and rating response order on the association between metacognitive ratings and identification task accuracy (Siedlecka et al., 2016; Wierzchoń et al., 2014). These studies’ authors suggested that carrying out a behavioral response acts as an additional source of evidence for metacognitive judgments. Following this, Anzulewicz et al. (2019) listed four possible mechanisms through which action planning or execution could influence reported awareness. They point to possible (1) indirect effects that stem from motor cortex activity that affects cognitive processing, (2) stimulus-related evidence accumulation being influenced by attentional engagement, (3) enhancement of performance monitoring, (4) integration of additional (including post-perceptual) evidence with the evidence accumulation process.

It has been shown that the evidence accumulation process is strongly coupled with the presence of perceptual stimulation, but it continues even after its disappearance – at least up to the moment at which one identifies the stimulus (Yeung & Summerfield, 2012) and might persist after decision to inform metacognitive judgments (Murphy et al., 2015). The hypothesis that post-perceptual information can concurrently influence metacognitive judgment is supported by Gajdos et al. (2019). The authors showed that higher confidence ratings were observed in trials in which an identification response to a stimulus was preceded by partial muscular activation. They argued that such muscle activity could contribute to participants’ confidence in their identification response to a stimulus, but it could not influence the identification task itself. However, these studies do not provide sufficient evidence to prove that partial muscular activations influence confidence judgment and not the opposite.

This issue of causal relation can be partially resolved by experimentally introducing additional M1 activity that is unrelated to the main task. In Siedlecka et al. (2019), this was achieved by including an irrelevant task that participants performed between stimulus presentation and PAS rating. Performing an additional motor response congruent with the response scheme of the identification task led to higher awareness ratings than when performing an incongruent one. At the same time, the congruence between the additional motor response and the identification task response was not related. Siedlecka et al.’s experiment provides arguments for the influence of motor system activity on visual awareness judgment. However, it cannot be ruled out that the additional response itself or the visual information from the additional task cue were responsible for the observed effect.

Assessment of the selective effect of motor information on visual awareness judgements requires directly altering motor cortex activity. Fleming et al. (2015) attempted this by applying single-pulse TMS either before or after a two-alternative forced-choice (2AFC) task followed by decision confidence rating. They showed that TMS applied to the dorsal premotor cortex (PMd) that was associated with the chosen response was associated with higher response confidence and consequently higher metacognitive efficiency (a quantitative measure of participants’ level of metacognitive ability, given a certain level of 2AFC task performance) than TMS associated with the unchosen response, while no evidence for the influence of TMS on identification accuracy was found. The TMS effect on mean confidence rating was observed for TMS applied both before and after the identification response, thus suggesting the contribution of post-decision processes to confidence in one’s identification decision. None of these effects was observed for TMS applied to M1. Fleming et al. suggested that PMd but not M1 activity contributes to confidence ratings. However, both TMS intensity and the number of participants taking part in the experiment were relatively low, thus encouraging the collection of more evidence on this matter.

Considering the limitations of the previous research, we investigated whether externally introduced motor-related information can be integrated into judgment of visual awareness. To achieve this, we delivered twitch-causing TMS to an M1 representation of the index finger involved in providing identification responses. Moreover, we used verbally reported PAS to separate the TMS and the identification response to minimize TMS-induced motor activity’s interference with the activity that resulted from the intentional identification task decision. Based on Fleming et al.’s (2015) results, we did not expect to observe the influence of TMS on identification decision performance, including its RT. Based on Siedlecka et al.’s (2019) findings, we expected to observe higher awareness ratings in the M1 condition compared to the control condition (TMS to the interhemispheric cleft). In addition, we calculated response-specific metacognitive efficiency measures. Since in our experiment, the scale response preceded the identification response and we asked for perceptual awareness judgments (not confidence judgments) in identification task decisions, we did not expect to observe any difference in metacognitive efficiency between M1 and the control TMS condition.

To actively monitor the precision of TMS delivery, we recorded MEP (motor-evoked potential) amplitudes on the response finger that was contralateral to the stimulation side. It has been established that imagined unilateral movements increase the excitability of contralateral M1 (Facchini et al., 2002; Fourkas et al., 2006; Jeannerod, 1995). Previous research on MEP has shown that its amplitude can reflect the level of M1 excitability (Fitzgerald et al., 2002). For these reasons, we expected M1 excitability to be influenced by the preparatory motor plan for the subsequent identification response proportionally to the accumulated evidence for the identification decision. This should lead to a correlation between MEP amplitudes and PAS ratings as well as a correlation between PAS ratings and their response times (RTs), thus representing accumulated evidence for visual awareness judgment.

## 2. Materials and methods

The experiment was carried out in the TMS Laboratory at the Neurology Clinic of Jagiellonian University Hospital. The study was approved by the Ethics Committee of the Institute of Psychology at Jagiellonian University and was carried out in accordance with the guidelines for TMS research (Rossi et al., 2009; Rossini et al., 2015) and the Declaration of Helsinki (Holm, 2019).

### 2.1 Participants

Healthy volunteers meeting the criteria for participation in TMS studies (no history of neurological disorders, psychiatric disorders, head injury etc., as assessed by a safety screening questionnaire) and with normal or corrected-to-normal vision were recruited using advertisements on social media. One participant dropped out due to TMS-induced headache, while 46 participants (2 left-handed, 11 males, M_age_ = 23.2, range = 19-37) completed the study. The general purpose of the experiment was explained to participants, and they were informed that they could withdraw at any time without giving a reason. Prior to the experiment, the participants completed safety screening questionnaires and signed informed consent forms. After the experiment, they received monetary compensation (160 PLN).

### 2.2 Session overview

The experiment was conducted using a within-participant design in a single session. Participants practised (15 trials, ~2 min) the procedure, with the identification task preceding the PAS rating within each trial (Ramsøy & Overgaard, 2004; Sandberg et al., 2010). Then a 1-up-3-down staircase was used to estimate the stimulus contrast (100 trials; step size 0.5%, starting with 9% of the maximal stimulus contrast of the monitor) that would lead to approx. 79% correct responses. The median contrast for each PAS rating was calculated based on all trials acquired in the staircase procedure (~5 min) for use in the following experimental procedure, in which four fixed contrasts were used in a random manner and with equal probability. The used contrasts did not differ between experimental conditions.

Subsequently, individual resting motor thresholds (RMTs) for TMS were determined, and participants completed a 32-trial training session that was identical to the experimental procedure: TMS pulses were applied to the left M1, and the PAS rating was followed by the identification task response. Finally, they completed the experimental task, which consisted of two conditions in four counterbalanced blocks (two blocks of TMS to M1, and two blocks of TMS to interhemispheric cleft, alternately). Each block consisted of 100 trials, which summed up to 400 trials that took about 45 minutes to complete.

### 2.3 Stimuli and procedure

The task was coded in PsychoPy software (Peirce, 2007) and was run on a PC. Participants placed their heads on a chinrest, 60 cm away from an LCD monitor (1920 × 1080 resolution, 60 Hz refresh rate). A microphone was attached to the chinrest for the purpose of PAS verbal responses recording. Stimuli consisted of Gabor patches tilted left or right (−45 or 45 degrees of rotation from vertical angle, respectively; 128 × 128 pixels, which translated to 3.2 degrees of visual angle) presented centrally on the screen. A white noise patch was presented with the stimulus to reduce its visibility. Fig. 1 outlines the temporal organization of an experimental trial and provides the methodological details of the procedure.

**Fig. 1.**
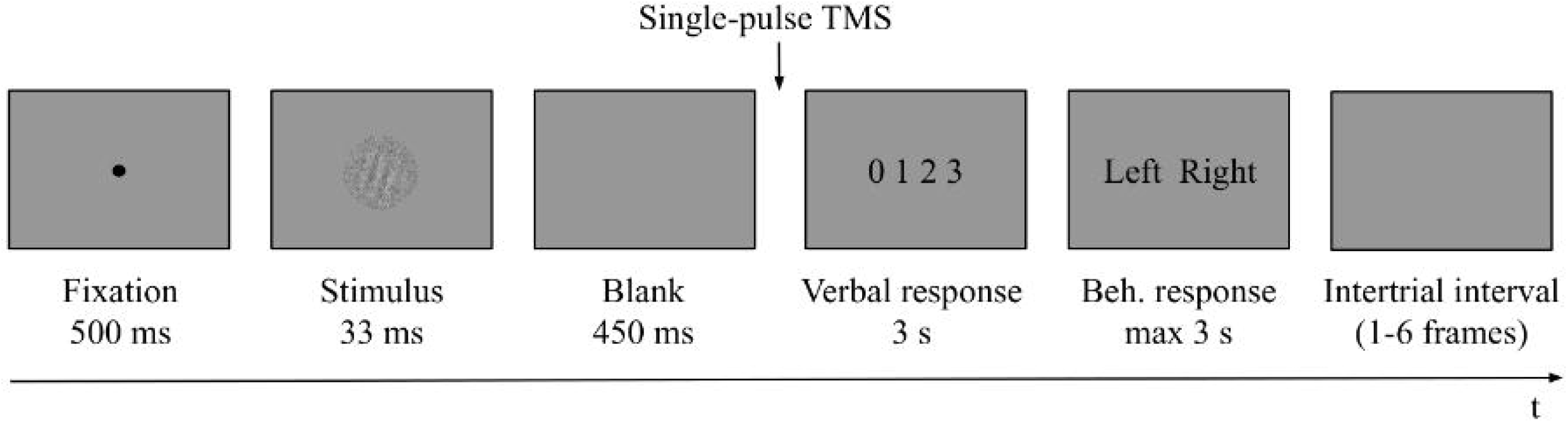
Schematic illustration of an experimental trial. First, a fixation dot was presented for 500 ms. A Gabor patch masked with white noise was then displayed for 33 ms, followed by an empty screen for 450 ms. Subsequently (i.e. 483 ms from the stimulus onset), a TMS pulse was administered and a screen prompting the PAS rating (with the points of the scale defined as 0 = no experience; 1 = a brief glimpse; 2 = an almost clear experience; 3 = a clear experience) was displayed for 3 seconds dedicated to provide a verbal response. Irrespective of whether a verbal response was provided or not, PAS was followed by a screen prompting a behavioral response to the identification task that was displayed for 3 seconds or until a keyboard button press was made (either “Z” with the left index finger or “M” with the right one). Participants did not receive any feedback about their performance. Trials were separated with intertrial intervals of variable length.

### 2.4 TMS parameters

Biphasic TMS was delivered with a Magstim Super Rapid^2^ Plus^1^ stimulator using a 70 mm Double Air Film Coil at 110% of the individual RMT (average intensity = 65.87% of the maximal stimulator output; MSO, standard deviation; SD = 10.67). The electromyographic signal was recorded from the first dorsal interosseous (FDI) muscle of the right index finger throughout the whole experimental procedure. The individual RMT estimation started with applying TMS at 50% of MSO to the left M1. Then, by varying the stimulation intensity, the site where suprathreshold TMS induced the maximal twitch in the right index FDI muscle was established. Afterwards, the lowest intensity that resulted in an MEP of more than 50μV peak-to-peak amplitude in five out of ten consecutive trials was determined. In the control condition, TMS was applied to the interhemispheric cleft between the superior parietal lobules, with the coil handle pointing backwards. The site of stimulation and the tangential position of the coil in relation to the scalp were monitored using the average brain template in the Brainsight 2.3 neuronavigation system. For the M1 stimulation, the main axis of the coil was oriented at 45° offset from the posterior-anterior (PA) direction, but it remained untilted for the control condition. The current induced in the brain was PA-AP. Participants wore earplugs for noise protection throughout the duration of TMS.

### 2.5 Data analysis

No statistical analyses were conducted before the completion of the experiment and no participants who completed the experiment were excluded from the analysis. Trials with no PAS response and identification response were removed; the remaining data (17,969 trials, 97.7%) were analyzed using the R statistical environment (R Core Team, 2019). We used mixed-effects regression models fitted with the lme4 package (Bates, Mächler, Bolker, & Walker, 2015). To obtain approximate p-values via Satterthwaite’s method, we used the lmerTest package (Kuznetsova et al., 2017). Additionally, we used the phia (De Rosario-Martinez, 2015) and emmeans packages (Lenth, 2019) for pairwise comparisons, employing Tukey’s method for family-wise error rate correction. We used code provided in an implementation of response-specific meta-d’ (Maniscalco & Lau, 2014) to calculate (1) identification task sensitivity index *d’*, (2) type 1 criterion indicating identification response bias, and (3) M-ratio (meta-*d’*/*d’*), which is a measure of metacognitive efficiency in which metacognitive sensitivity (operationalized with meta-*d′*) is corrected for objective task sensitivity (operationalized with *d′*; Fleming & Lau, 2014). A M-ratio indicates the amount of evidence available for metacognitive judgement relative to the amount of evidence available for objective decision, e.g. the M-ratio value of 0.8 indicates that 20% of the sensory evidence available for the objective decision is lost when making metacognitive judgments, while M-ratio value of 1.2 suggests that more evidence is available for metacognitive judgements than for objective decision that can be due to further processing of stimulus information or gaining non-perceptual information (Fleming, 2017). For MEP-related calculations, for every trial, the highest peak-to-peak amplitude was determined in the 75 ms after the TMS pulse, irrespective of the condition. We intended to present results from the full dataset, therefore we did not limit analysis to trials in which the MEP amplitude exceeded 50 μV, as is commonly done (Anderson & George, 2009). In order to convert verbal recordings with PAS ratings into a machine-readable format, we used Python’s Speech Recognition package (Zhang, 2017); we calculated speech onset with Chronset and trials for which the algorithm failed were corrected manually. RTs were measured either from the PAS screen or the onset of the identification task response cue. We use congruence between TMS and identification response as a fixed factor. Although no TMS-induced movement was present in the control condition, we used congruence to refer to right-hand responses. Because TMS was limited to the left hemisphere, all responses provided with the right index finger were TMS-response congruent (congruent trials, n = 8,933), while all those provided with the left index finger were TMS-response incongruent (incongruent trials, n = 9,036). We used non-directional tests with *α* level set at 5%.

## 3. Results

### 3.1 Identification task

Identification task accuracy data were analyzed using a logistic mixed-effects regression model with condition and congruence as fixed effects. All effects were taken as random at the participant level. As expected, no significant differences in accuracy were found (see Table 1 for the model summary and Fig. 2): neither between the control and M1 conditions within congruent trials (*z* = 0.87, *p* = .384), nor between incongruent and congruent trials within the M1 condition (*z* = −0.07, *p* = .944). No interaction between condition and congruence was observed (*z* = −0.68, *p* = .498). Taken together, no evidence was thus found for a general effect of the M1 condition on the accuracy, despite the high number of trials and participants.

**Table 1.** Results summary of the mixed-effects logistic regression model for accuracy with TMS condition and TMS-response congruence as fixed effects; participant-specific condition effect, congruence effect, and intercept were used as random effects.

|  | Estimate | SE | z | p |
| --- | --- | --- | --- | --- |
| (Intercept) | 1.738 | 0.11 | 15.58 | < .001*** |
| TMS condition | 0.063 | 0.07 | 0.87 | .384 |
| TMS-response congruence | - 0.008 | 0.12 | -0.07 | .944 |
| TMS condition x TMS-response congruence | - 0.059 | 0.09 | -0.68 | .498 |

Signif. codes: < .001 '\*\*\*', .001 '\*\*', .01 '\*', .05 '.', .1

**Fig. 2.**
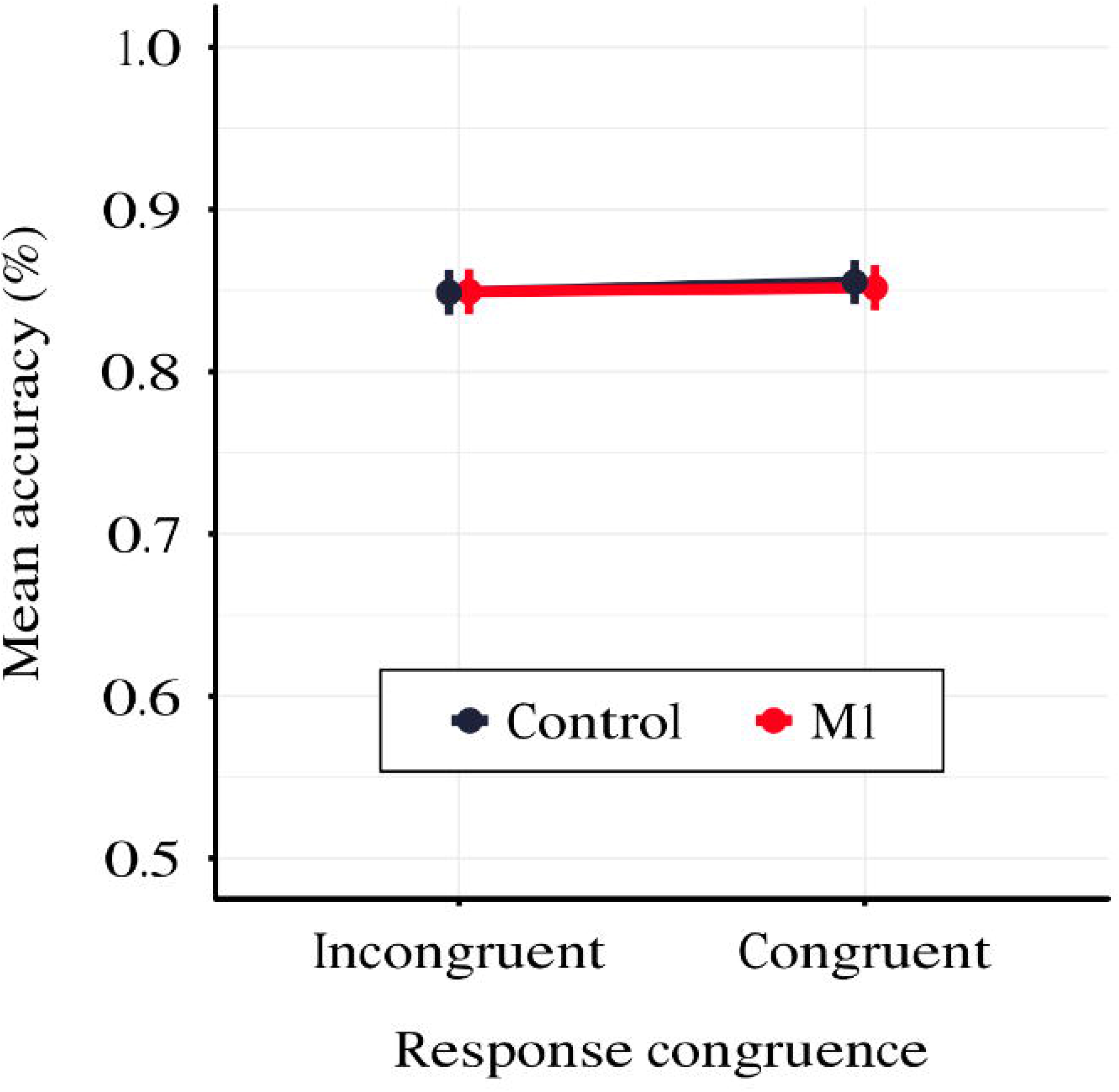
Mean identification task accuracy depending on TMS condition and TMS-response congruence. Error bars represent SEs.

Moreover, we calculated *d’* and the type 1 criterion for every participant for both TMS conditions separately; we fitted a linear mixed-effects model with condition as a fixed factor, and a participant-specific intercept. No difference in *d’* (*t*(45.0) = −0.86, *p* = .394) and the type 1 criterion (*t*(45.0) = −1.107, *p* = .274) between the M1 condition and the control condition was observed. The analysis thus did not find evidence for a difference in the identification ability and the response criterion in the identification task across conditions.

To investigate identification RTs, we fitted a linear mixed-effects regression model with interactions between condition, congruence, and PAS rating as fixed effects. All effects were taken as random at the participant level. We found that the RTs in the M1 condition were significantly longer than in the control condition within congruent trials (*t*(86.78) = 2.29, *p* = .024); also, in the M1 condition, congruent trials took longer than incongruent ones (*t*(99.99) = 3.05, *p* = .003). Additionally, we were interested in how these RT differences manifested across PAS ratings. Conditional pairwise comparisons revealed that identification RTs for the middle ratings were significantly longer in the M1 condition compared to the control condition within congruent trials. The same was observed for congruent trials as compared to incongruent trials within the M1 condition (see Table 2, Fig. 3). In sum, identification responses were slower when the TMS influenced the muscle activity of the hand with which participants responded to stimuli which were detected but not seen clearly, thus indicating an extended evaluation process in these cases.

**Table 2.**
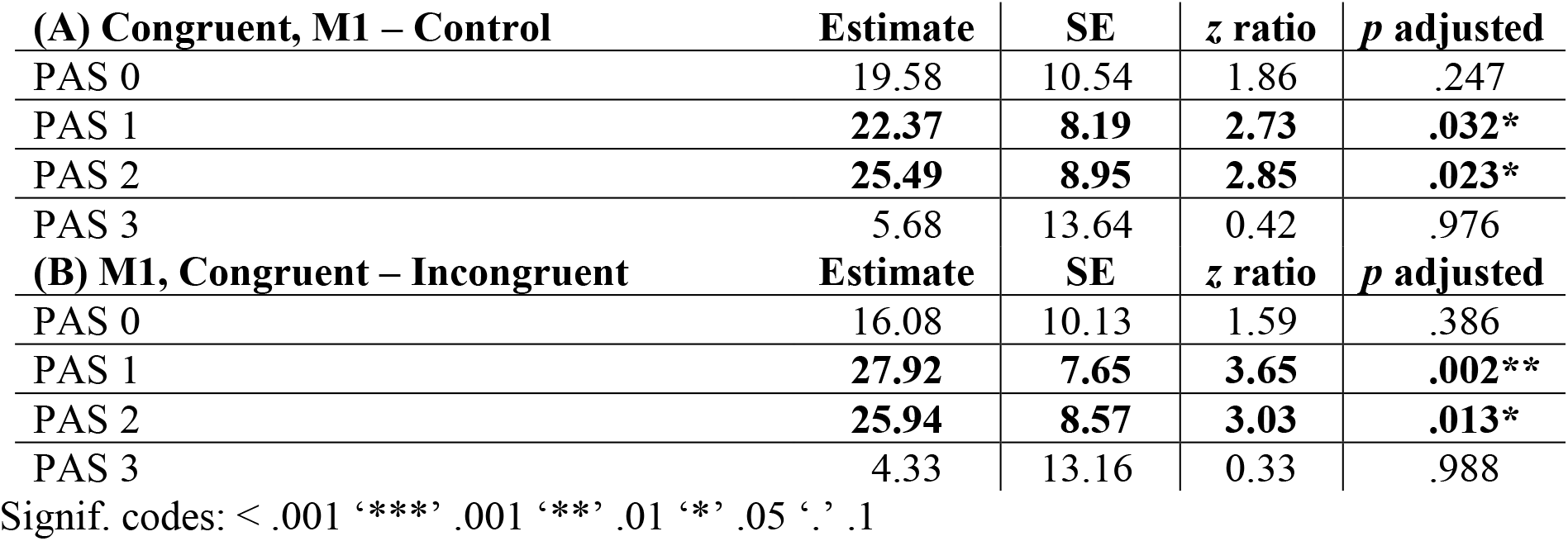
Pairwise comparisons of identification RTs’ regression coefficients for the linear mixed-effects regression model with interactions between condition, congruence and PAS rating as fixed effects, and with participant-specific condition effect, congruence effect, and intercept as random effects. P values adjusted with Tukey correction method. (A) Comparisons of estimates for each PAS rating between M1 and control condition in congruent trials. (B) Comparisons of estimates for each PAS rating between congruent and incongruent trials in the M1 condition.

| **(A) Congruent, M1 – Control** | **Estimate** | **SE** | **z ratio** | **p adjusted** |
| --- | --- | --- | --- | --- |
| PAS 0 | 19.58 | 10.54 | 1.86 | .247 |
| PAS 1 | **22.37** | **8.19** | **2.73** | **.032\*** |
| PAS 2 | **25.49** | **8.95** | **2.85** | **.023\*** |
| PAS 3 | 5.68 | 13.64 | 0.42 | .976 |
| **(B) M1, Congruent – Incongruent** | **Estimate** | **SE** | **z ratio** | **p adjusted** |
| PAS 0 | 16.08 | 10.13 | 1.59 | .386 |
| PAS 1 | **27.92** | **7.65** | **3.65** | **.002\*\*** |
| PAS 2 | **25.94** | **8.57** | **3.03** | **.013\*** |
| PAS 3 | 4.33 | 13.16 | 0.33 | .988 |

Signif. codes: < .001 '\*\*\*' .001 '\*\*' .01 '\*' .05 '.' .1

**Fig. 3.**
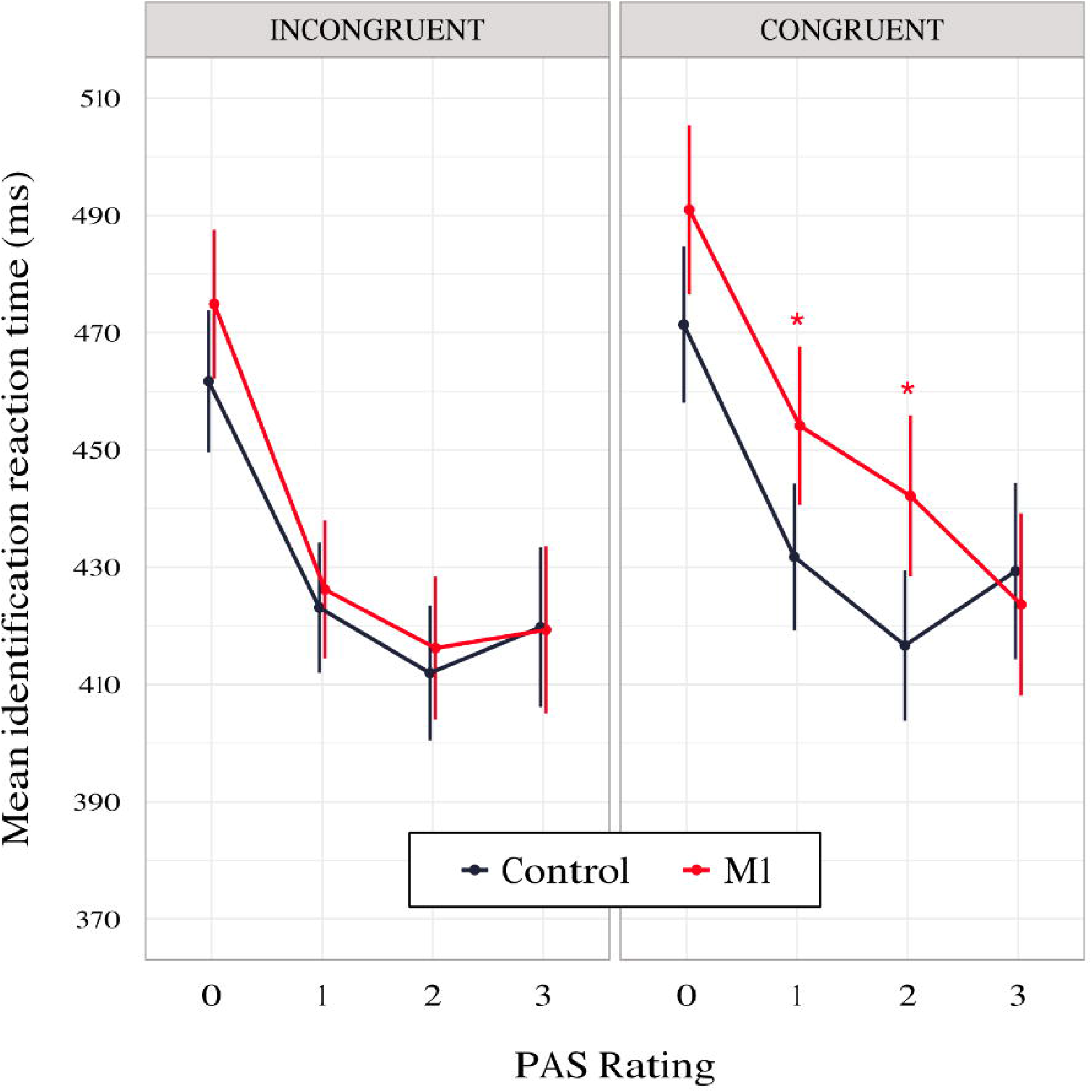
Mean identification task RT depending on TMS-response congruence, TMS condition and PAS rating. Error bars represent SEs.

### 3.2 PAS ratings

To test the impact of TMS on PAS ratings, we fitted a linear mixed-effects model with interaction between condition and congruence as fixed effects. All effects were taken as random at the participant level. We used linear modelling to achieve comparability with the previous study of Fleming et al. (2015) and because the available implementations of ordinal models do not allow random effects in individual thresholds (Bürkner and Vuorre, 2019). We observed a significant interaction between condition and congruence (*t*(17829.20) = −2.30, *p* = .021, see Table 3 for regression model summary). We reparameterized the model to check the effect of the TMS condition that was nested in the TMS-response congruence effect and observed a numerically higher mean PAS rating in M1 compared to the control condition in congruent (*t*(66.46) = 1.76, *p* = .083) but not incongruent trials (*t*(65.84) = −0.16, *p* = .876).

**Table 3.** Results summary of the linear mixed-effects model for PAS ratings with condition and TMS-response congruence as fixed effects; participant-specific condition effect, congruence effect, and intercept were used as random effects.

|  | **Estimate** | **SE** | **t (df)** | **p** |
| --- | --- | --- | --- | --- |
| (Intercept) | 1.344 | 0.074 | 18.20 (45.7) | < 0.001*** |
| TMS condition | - 0.050 | 0.028 | - 1.76 (66.5) | .083. |
| TMS-response congruence | - 0.057 | 0.043 | - 1.34 (52.6) | .185 |
| TMS condition x TMS-response congruence | **- 0.054** | **0.024** | **- 2.30 (17829.3)** | **.021\*** |

Signif. codes: < .001 '\*\*\*' .001 '\*\*' .01 '\*' .05 '.' .1

Since Fleming et al. (2015) observed a similar effect in correct trials and a reversed pattern (higher confidence in incongruent than congruent trials) in incorrect trials, we ran the model separately for subsets of correct (n = 14,841) and incorrect (n = 3,128) identification response trials. The results pattern did not depend on accuracy. For correct trials, we observed a significant effect of interaction between condition and congruence (*t*(14,751.0) = −2.54, *p* = .011) and a significantly higher mean PAS rating in M1 compared to the control condition incongruent (*t*(70.93) = 1.20, *p* = .050) but not in incongruent trials (*t*(70.33) = 0.16, *p* = .795; Fig. 4, panel A and C). In incorrect trials, a significant interaction between condition and congruence (*t*(1,650.52) = −2.01, *p* = .044) was also present. There was a significantly higher mean PAS rating in M1 compared to the control condition within congruent (*t*(90.90) = 2.04, *p* = .044) but not incongruent trials (*t*(88.23) = 0.44, *p* = .659; Fig. 4, panel B and D). The reparameterization of the model did not show an effect of congruence (*t*(50.71) = 0.94, *p* = .354) in M1 correct trials, but it revealed a significant difference between congruent and incongruent trials (*t*(74.70) = 2.05, *p* = .044) in incorrect M1 trials (see Fig. 4, panel A and B).

**Fig. 4.**
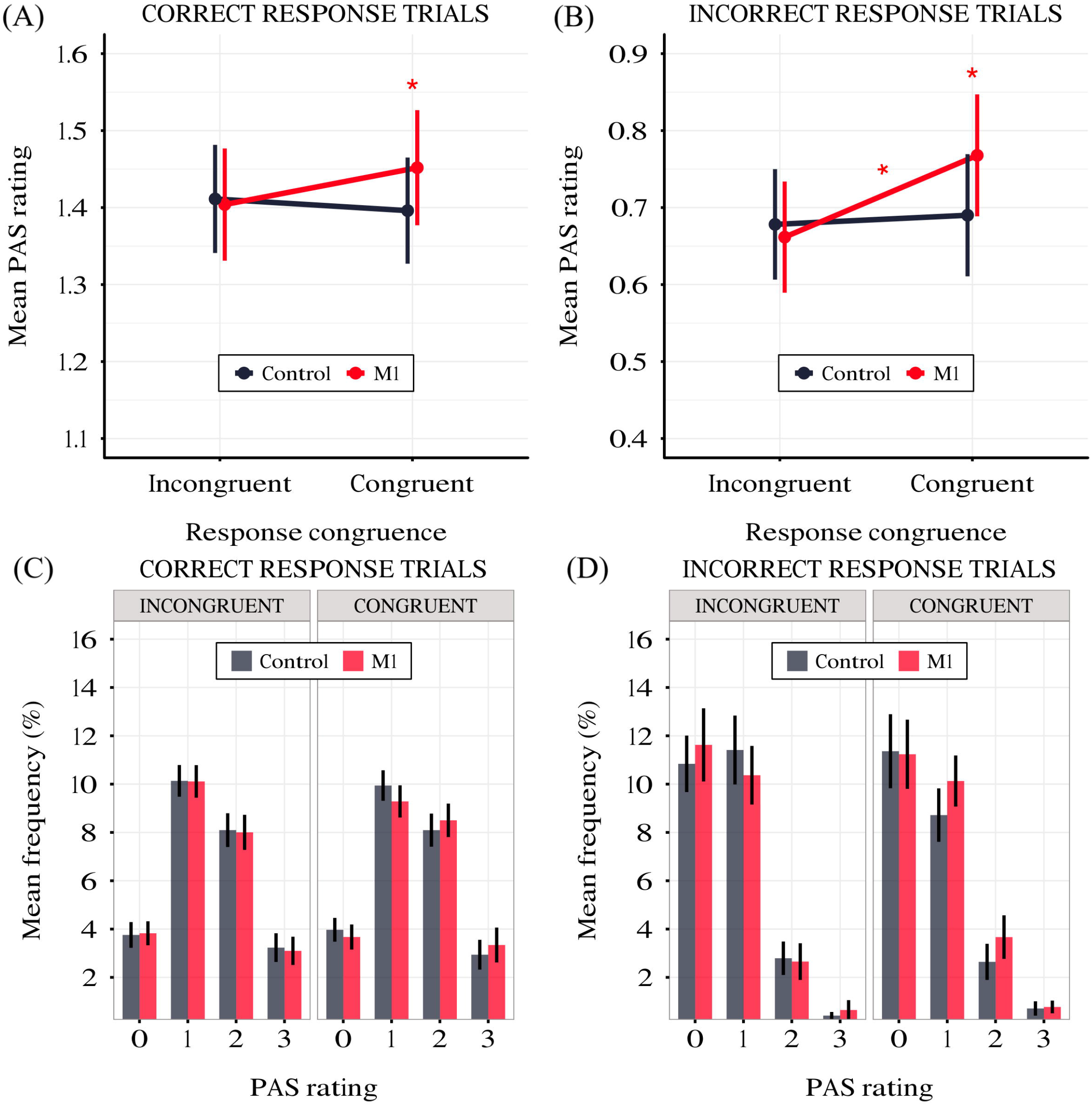
(A) Mean PAS rating depending on TMS condition and TMS-response congruence in correct trials. (B) Mean PAS ratings for TMS condition and TMS-response congruence in incorrect trials. (C) PAS ratings’ distributions depending on TMS-response congruence and TMS condition in correct trials. (D) PAS ratings’ distributions depending on TMS-response congruence and TMS condition in incorrect trials. Error bars represent SEs.

Additionally, to compare RTs of PAS ratings, we fitted a mixed-effects linear regression model with interactions between condition, congruence, and PAS ratings as fixed effects. All effects were taken as random at the participant level. This analysis revealed that the PAS rating RTs in the M1 condition were significantly longer than in the control condition (*t*(59.38) = 3.58, *p* < .001) within congruent trials. Since no interaction between condition and congruence was observed (*t*(17,749.88) = 0.31, *p* = .754), the effect was observed for both congruent and incongruent trials. The pairwise comparisons revealed evidence that the effect applied to the two lowest ratings’ RTs (see Table 4 and Fig. 5 for pairwise comparisons).

**Table 4.**
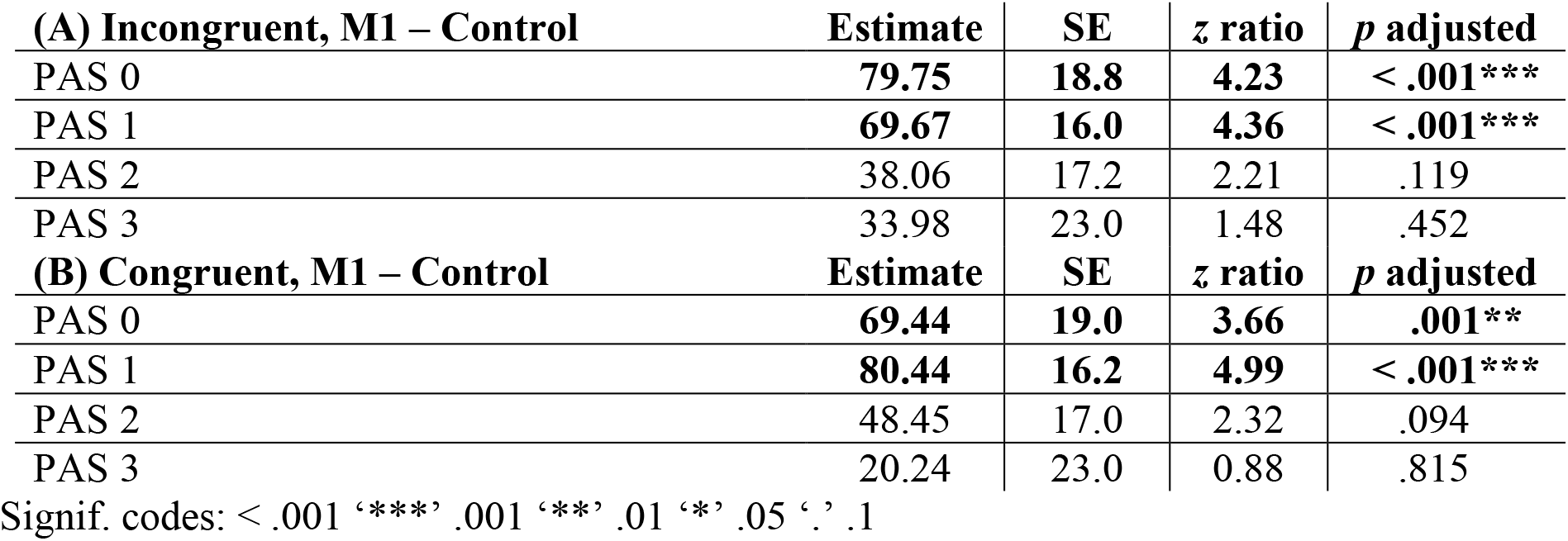
Pairwise comparisons of PAS RTs’ regression coefficients of the linear mixed-effects regression model with interactions between condition, congruence and PAS ratings as fixed effects, and participant-specific condition effect, congruence effect, PAS rating effect, and intercept as random effects. P values adjusted with Tukey correction method. (A) Comparisons of estimates for each PAS rating between M1 and control condition in incongruent trials. (B) Comparisons of estimates for each PAS rating between M1 and control condition in congruent trials.

**Fig. 5.**
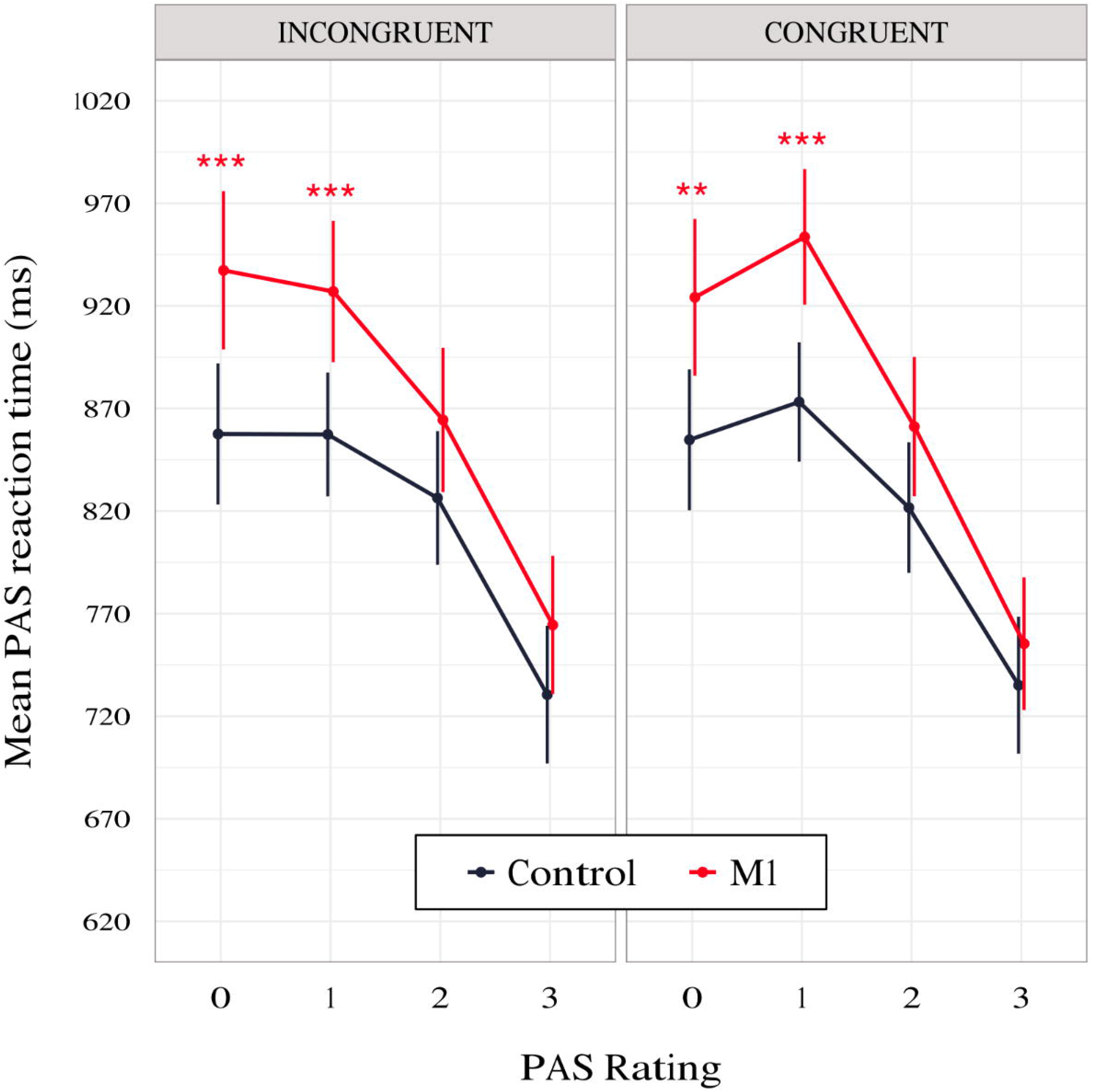
PAS ratings’ mean RTs as a function of response congruence and TMS condition. The error bars represent SEs.

### 3.3 M-ratio

To test whether there was a difference in metacognitive efficiency between the M1 and control conditions, we calculated M-ratios for every participant for TMS and congruence conditions separately. We fitted a linear mixed-effects model with condition and congruence as fixed factors, and with participant-specific congruence effect and intercept as random effects. We found no significant effect of condition (*t*(91.6) = −0.15, *p* = .882) or congruence (*t*(90.0) = 0.98, *p* = .332), and no interaction between condition and congruence was observed (*t*(90.0) = −0.48, *p* = .630; see Fig. 6). It should be noted that due to the greater analysis complexity, these tests may have lower statistical power than the other analyses presented in the paper (Kristensen et al., 2020). However, because there was an increase in PAS ratings for both correct and incorrect trials, we did not expect to observe a difference in M-ratio.

**Fig. 6.**
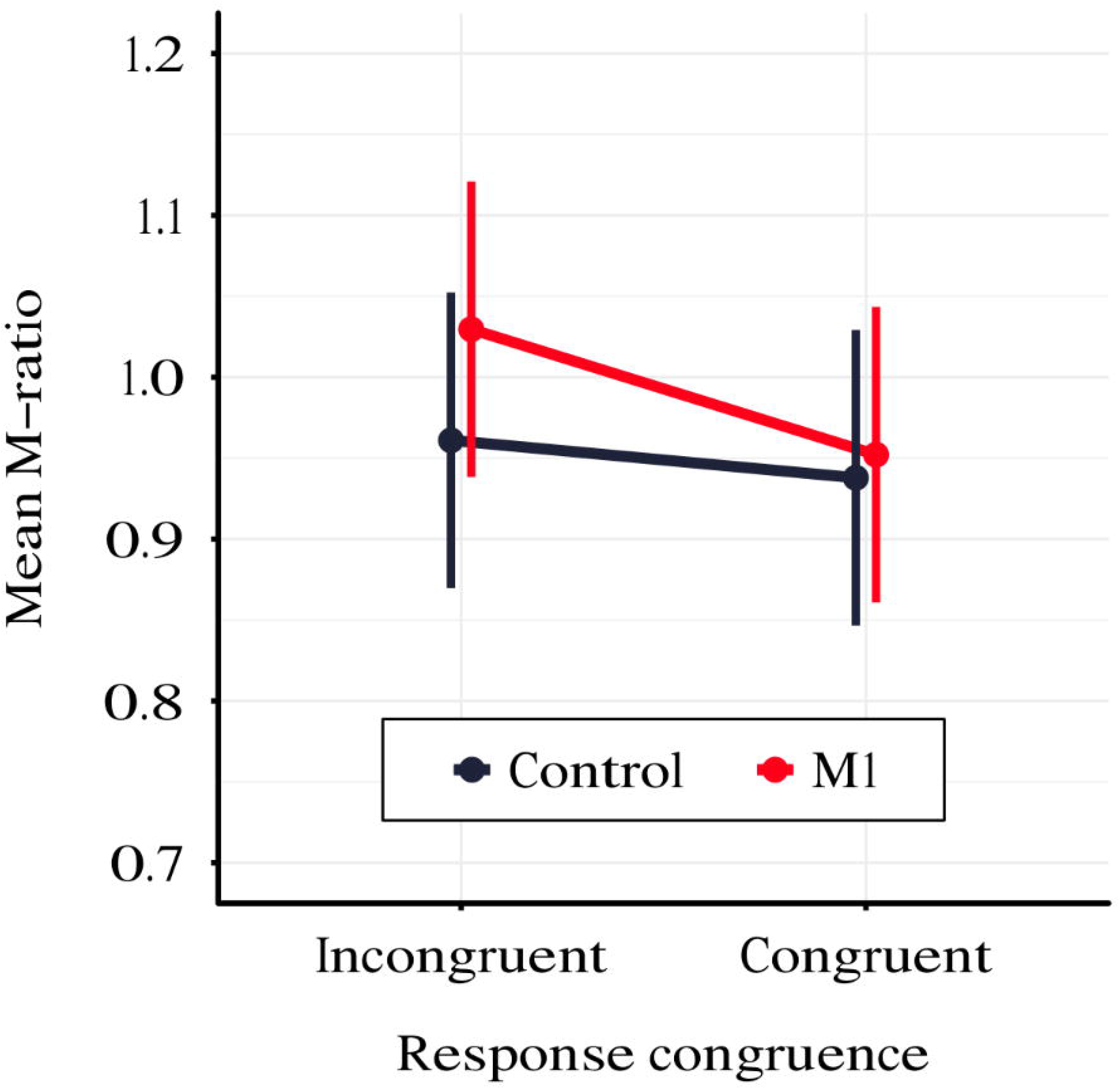
Mean M-ratio depending on TMS condition and TMS-response congruence. The error bars represent SEs.

### 3.4 MEP amplitudes

To test differences in MEP amplitudes, we fitted a linear mixed-effects model with interaction between condition, congruence and PAS rating as fixed effects, and with participant-specific condition effects, congruence effect, and intercept as random effects. Since only M1 TMS was supposed to influence the motor cortex, a significant difference between MEP amplitudes in the M1 condition compared to the control condition was observed (*t*(45.81) = 10.61, *p* < .001). Interestingly, there was a significant interaction between congruence and condition (*t*(17465.41) = 5.70, *p* < .001), and the results of the model reparameterization showed significantly higher MEP amplitudes in congruent trials (*t*(116.09) = 6.55, *p* < .001). Additionally, we were interested in determining whether this difference was related to PAS ratings. Therefore, we performed pairwise comparisons of MEP amplitudes for each PAS rating between congruent and incongruent trials in the M1 condition. Comparing amplitudes of MEPs across PAS ratings gradually yielded significant differences (see Table 5 and Fig. 7 for detailed results).

**Table 5.** Pairwise comparisons of mean MEP amplitude regression coefficients of the linear mixed-effects regression model with interactions between condition, congruence and PAS ratings as fixed effects, and with participant-specific condition effect, congruence effect, and intercept as random effects. P values adjusted with Tukey correction method. Comparisons of estimates for each PAS rating between congruent and incongruent trials in the M1 condition.

| **M1, Congruent – Incongruent** | **Estimate** | **SE** | **z ratio** | **p adjusted** |
| --- | --- | --- | --- | --- |
| PAS 0 | **44.46** | **16.16** | **2.75** | **.012\*** |
| PAS 1 | **33.46** | **11.97** | **2.80** | **.012\*** |
| PAS 2 | **50.43** | **13.56** | **3.72** | **< .001\*\*\*** |
| PAS 3 | **114.23** | **21.20** | **5.39** | **< .001\*\*\*** |

Signif. codes: < .001 '\*\*\*' .001 '\*\*' .01 '\*' .05 '.' .1

**Fig. 7.**
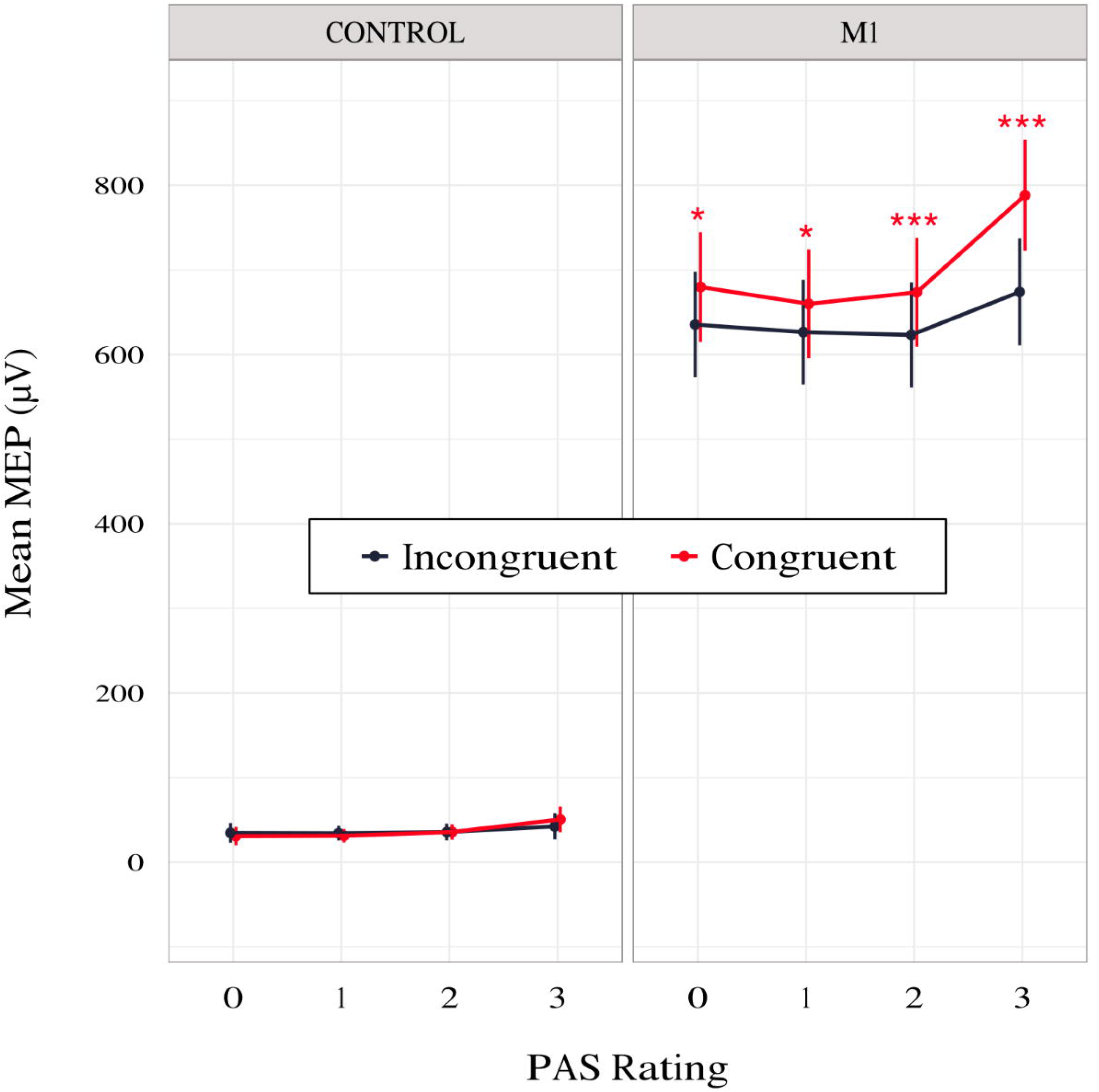
Mean amplitudes of MEPs for each PAS rating depending on TMS condition and response congruence. Error bars represent SEs.

To determine if it is possible to discern PAS rating from the preceding MEP, we compared amplitudes from trials with different PAS ratings. Only trials with rating 3 (a clear experience) were significantly different from the others, irrespective of congruence (see Table 6 for all pairwise comparisons).

**Table 6.** Pairwise comparisons of mean MEP amplitude regression coefficients of the linear mixed-effects regression model with interactions between condition, congruence and PAS ratings as fixed effects, and with participant-specific condition effect, congruence effect, and intercept as random effects. P values adjusted with Tukey correction method. Comparisons of estimates between PAS ratings for congruent and incongruent trials in the M1 condition.

| **M1** | **PAS** | **Estimate** | **SE** | **z ratio** | **p adjusted** |
| --- | --- | --- | --- | --- | --- |
| Incongruent | 0 - 1 | 8.94 | 13.03 | 0.68 | .902 |
|  | 0 - 2 | 12.20 | 14.49 | 0.84 | .834 |
|  | 0 - 3 | - 38.64 | 19.19 | - 2.01 | .183 |
|  | 1 - 2 | 3.26 | 11.83 | 0.29 | .993 |
|  | 1 - 3 | **- 47.58** | **17.39** | **- 2.74** | **.031\*** |
|  | 2 - 3 | **- 50.85** | **17.53** | **- 2.90** | **.020\*** |
| Congruent | 0 - 1 | 19.94 | 13.60 | 1.47 | .458 |
|  | 0 - 2 | 6.23 | 14.62 | 0.43 | .973 |
|  | 0 - 3 | **- 108.41** | **19.16** | **- 5.66** | **< .001\*\*\*** |
|  | 1 - 2 | - 13.71 | 11.76 | - 1.17 | .648 |
|  | 1 - 3 | **- 128.35** | **17.11** | **- 7.50** | **< .001\*\*\*** |
|  | 2 - 3 | **- 114.63** | **17.05** | **- 6.72** | **< .001\*\*\*** |

Signif. codes: < .001 '\*\*\*' .001 '\*\*' .01 '\*' .05 '.' .1

## 4. Discussion

In the present study, we aimed to gain insight into the influence of the motor system on perceptual awareness judgments. We determined whether TMS-induced activity that was delivered to M1 following stimulus presentation altered participants’ judgments of stimulus awareness, as indexed by PAS ratings. Our results show that TMS congruent to participants’ responses increased the reported stimulus awareness, but there was no evidence for altering the extent to which they are objectively sensitive to visual information. Moreover, no identification task bias was observed. We observed longer awareness rating RTs in the M1 condition in trials accompanied by stimulus awareness that was vaguer than an almost clear experience. Despite using a procedure with a delayed identification task, we observed longer identification RTs in the M1 condition, in which the TMS effect on RTs was limited to TMS-response-congruent trials. Additionally, we attempted to determine whether TMS had an influence on the metacognitive efficiency measure, but we found no evidence to support such a claim. Finally, MEP amplitudes were related to PAS ratings and response congruence.

Based on the presented data, we suggest that the externally induced activity in M1 served as additional non-diagnostic evidence for the evidence-accumulation processes underlying visual awareness judgment and stimulus identification decisions. In response to TMS, we observed prolonged activity of these processes, predominantly when the perceptual evidence was not fully decisive. Moreover, the process of stimulus-related evidence accumulation also seems to be reflected in M1 excitability, which is implicated by MEP amplitude.

### 4.1 Motor information influences visual awareness judgments

Our primary goal was to determine whether the activity of the motor system can contribute to perceptual judgments. The work of Siedlecka et al. (2019) has already shown that irrelevant motor responses that share a response scheme with a visual stimulus identification task increase reported stimulus awareness. However, their design did not allow the exclusion of confounding factors such as the introduction of additional visual information or attentional engagement, because participants were explicitly instructed and cued to perform an additional response (but see: Siedlecka, Paulewicz, Koculak, 2020). In this study, we expanded upon their paradigm by applying TMS to M1 to reduce the influence of these confounds. Nevertheless, our conclusions reinforce those of Siedlecka and colleagues (2019): additional motor system activity can be incorporated in perceptual awareness judgment.

Unlike Siedlecka et al. (2019), we only found indications for additional evidence integration in response congruent trials, whereas their results showed an increase in PAS ratings irrespective of response congruence. This could be a consequence of the fact that participants in their study performed an intentional response before providing a rating. The additional task was very simple so that participants could have both motor plans prepared in each trial before responding to a visual cue, which would increase motor cortex activity in both hemispheres. In our study, TMS-induced motor activity could be weaker than that related to actual movement, which might explain its specific effect. Our results provide confirmatory evidence that the findings reported by Siedlecka, et al. (2019) were a result of motor activity.

A similar approach was undertaken by Fleming et al. (2015), who used single-pulse TMS either before or immediately after the 2AFC task response. In separate experiments, TMS was applied to either PMd or M1. The results of PMd TMS revealed higher confidence in TMS-response-incongruent (as compared to congruent) response trials in error trials within the pre-response TMS condition. Moreover, for correct trials in the pre-response TMS condition, a tendency towards lower confidence in trials with TMS-incongruent as compared to TMS-congruent responses was observed. This difference was significant in the post-response TMS condition, but no M1 TMS effect was observed. An explanation of the discrepancies between their and our M1 TMS results could be the different timing of TMS, its intensity, or the substantial difference in sample size (23 vs. 46). Our conclusions suggest that M1 TMS might influence metacognitive judgment (although not necessarily metacognitive sensitivity). Besides, it might also suggest that the PMd TMS effect observed in Fleming’s study could actually have partly been a result of an increase in confidence in TMS-response-congruent trials, as indicated by their results, especially in the post-response TMS condition. Providing additional activity to the motor cortex might strengthen ongoing activity and increase confidence in an already chosen response. In this interpretation, PMd activity would reinforce the motor plan related to the TMS congruent response that would compete with the plan in the hemisphere related to the TMS incongruent response. Thus, not only activity in M1 congruent to TMS would increase, but this increase would cause inhibition of M1 in the other hemisphere (Reis et al., 2008); as a consequence, activity in M1 would not only increase confidence in congruent responses but would also decrease it in incongruent ones.

In addition, as might be supported by the analyses of metacognitive efficiency measures, PMd and M1 might differently impact metacognitive processes. Fleming et al. (2015) reported lower metacognitive efficiency (measured with M-ratio) in incongruent trials. Specifically, this was observed only for the PMd TMS that was delivered prior to the identification response. These results might further support the assumption that information related to PMd activity is incorporated into the evaluation of action performance. This could happen through increased activity of M1 related to the alternative motor scheme, which represents evidence against the chosen response. The facilitation of the alternative response could impact confidence and consequently lead to decreased M-ratio in TMS-response-incongruent trials. This seems in line with research showing temporarily increased excitability in M1 in response to PMd stimulation (Koch et al., 2006). Importantly, stimulation of the left PMd is often reported as affecting M1 in both hemispheres (Fujiyama et al., 2016). In our study, additional M1 activity would increase the amount of evidence for the congruent response (correct or not), resulting in no change in metacognitive efficiency measures while still impacting overall metacognitive ratings.

In the study of Fleming et al. (2015), the only RT effect observed was related to longer discrimination task and confidence rating RTs in the PMd and M1 conditions when TMS was applied after the discrimination response as compared to before it. This effect was not replicated in the second experiment reported in their paper. In comparison, our results show longer PAS RTs in M1 when stimulus awareness was absent or unclear. We consider these results to be a consequence of an evidence-accumulation process, but this process is no longer required in trials with high stimulus awareness because no additional evidence is needed. When the stimuli experience is not clear, this process operates for longer and/or with a lesser amount of evidence. This not only makes TMS-induced activity appear to be incorporated “on time” in a particular decision or judgment but also additional evidence can be of higher importance. It might be the case that this was possible partially due to the inclusion of the identification response at the end of the trial so the accumulation processes could last longer.

The nature of TMS experiments often makes the manipulation apparent to participants. In our study, due to finger movements, it might have been clear to them which condition was the experimental one, thus triggering observer-expectancy effects. However, this should result in differences between conditions in identification accuracy or increase PAS ratings for both congruent and incongruent trials, but these effects were not present in the data. For the observer-expectancy effect to be the case, participants would have to hold a specific belief about the experimenters’ expectation of higher awareness in the M1 condition and TMS-response congruent trials. They would have to remember to rate awareness as higher only when TMS is congruent with the identification task response, or when the Gabor is tilted to the right. The first would require the relatively difficult task of making a comparison with a response that follows an awareness rating. The second can be ruled out because PAS ratings for right-oriented Gabor stimuli provided with the left hand (incorrect trials in incongruent condition) were not higher as compared to the control. An alternative possible interpretations of our results are that TMS in TMS-response congruent condition triggered a distraction leading to attentional capture or influenced participants expectations, both resulting in that the participants paid greater attention to right-oriented Gabors in the M1 condition. If any of these was the case, then we should have observed a difference in the identification task performance and/or bias between the experimental conditions, but this was not the case. Therefore, although the design of the study cannot fully rule out attention or expectation effects, in our view their influence in this study is negligible.

### 4.2 Possible mechanisms of additional evidence integration

Our experiment provides evidence for a distinct path in a complex system that integrates information between perception, metacognition, and action. However, the structure of this system and the nature of the interactions between its parts is still largely unknown. This leaves an open question about the neuronal mechanism that leads to the effects observed in this study.

There is a growing body of evidence which shows that the prefrontal cortex (PFC), especially the dorsolateral prefrontal cortex (dlPFC), can be considered a key structure in integrating the information that is necessary for metacognitive judgments (Fleming & Dolan, 2012; Rounis et al., 2010). Assuming the central role of the dlPFC in awareness judgments, there are strong presumptions to treat the TMS effects observed in our study as an indirect influence because there are no direct connections between M1 and dlPFC (Passingham, 1993). This implies that the most probable route for the integration of information from M1 is through the primary somatosensory cortex (S1). Nevertheless, there is ample evidence for the reciprocal connections between M1 and S1 (Gandolla et al., 2014), through which M1 activity would influence S1 activity. Such S1 influence could resemble feedback information about the muscle activation of the response finger.

Information transfer between M1 and S1 through muscle activation could explain why our experiment, in which TMS intensity was above participants’ RMT, and the study by Siedlecka et al. (2019), in which participants performed an additional behavioral response, resulted in a significant influence of experimental manipulation on awareness ratings. Both explanations seem consistent with the results of Fleming et al. (2015), who used TMS intensity that was below the threshold of overt motor activity and thus limited the possibility of 1) sufficient direct influence from M1 on S1, or 2) sufficient muscle activity to cause somatosensory feedback. Similarly, Gajdos et al. (2019) suggest that pre-response partial muscle activation alters the somatosensory readout, which is later integrated into metacognitive judgment.

However, the posterior parietal cortex (PPC) would also likely be involved in integrating somatosensory information with perceptual evidence from other modalities. In normal circumstances, information from the sensorimotor feedback loop would be used to compare executed behavior with the motor plan that requires the engagement of frontal areas (e.g. dlPFC and PM). The more pronounced the mismatch, the lower one’s confidence in the accuracy of one’s action would be. This might be why the procedural manipulation of Fleming et al. (2015) resulted in a difference in metacognitive efficiency in the pre-response TMS condition. Their stimulation of PMd possibly altered the 2AFC task response execution, thus creating a mismatch that was caught by the error monitoring processes. However, TMS in our experiment was delivered early enough before the identification response to be integrated as additional evidence before a motor plan was fully formed. This would selectively increase the evidence for a stimulus associated with a particular motor plan, thus allowing participants to give higher metacognitive ratings in TMS-response-congruent trials. Crucially, early integration of this motor information would not create a mismatch between the planned and the performed response, so it did not lead to a change in metacognitive efficiency. The observation that TMS-related evidence interplayed with the selected motor plan suggests that either higher PAS ratings and longer identification RTs in M1 TMS have a common cause, or PAS response provides additional evidence for identification task decisions.

### 4.3 MEP as a measure of accumulated evidence

Our additional hypotheses concerned the possibility of using the MEP to quantify the neuronal correlate of perceptual evidence accumulation. MEP amplitude is frequently used as a read-out of M1 excitability state (Bestmann & Krakauer, 2015). Cognitive manipulation of spatial attention (Mars et al., 2007), values assigned to different responses (Klein-Flügge & Bestmann, 2012), or contextual uncertainty (Bestmann et al., 2008) can all influence M1 excitability. Crucially, M1 can be treated as a recipient of a decision process initiated in other brain areas that modulates its excitability (Klein-Flügge & Bestmann, 2012; Klein-Flügge et al., 2013). When a relation with a particular response is present, MEP amplitudes for chosen versus unchosen actions distinguish the forthcoming choice before completion of the decision process (Klein-Flügge & Bestmann, 2012).

Our electromyographic results go along with these findings. We found an effect of M1 TMS-response congruence: congruent trials were characterized by higher MEP amplitudes. This effect was observed predominantly when participants reported high stimulus awareness. These results seem to be complementary to the dynamics of evidence accumulation reflected in identification RTs. Taken together, they suggest that for clearly visible stimuli, when the necessary evidence has already been accumulated, the motor plan has been selected prior to TMS, thus increasing M1 excitability in preparation for execution of the response. Alternatively, no motor decision has been made, but the perceptual information about the stimulus has been passed from the visual cortex to M1, bypassing the PFC (Goodale, 2011). Presumably, within such conditions, additional evidence from the TMS does not play a crucial role in awareness judgment. Contrarily, while stimulus awareness is low, accumulation of evidence is still ongoing, thus allowing TMS to play a noticeable role.

Finally, changes in MEP amplitudes might reflect accumulation of stimulus-related evidence since the TMS-induced movement and the identification response were separated by several seconds, long before motor response execution. This seems possible based on the presence of connections from PPC to M1 (Gharbawie et al., 2011). There is a substantive body of evidence that PPC serves a multisensory integration function (Goodale and Miller, 1992; Kaas & Stepniewska, 2016; Koch et al., 2007) and plays an important role in performing voluntary movements, especially if they require visual input (Vingerhoets, 2014). In recent years there has been growing evidence that PPC has direct reciprocal connections to M1 (Isayama et al., 2019; Schulz et al., 2015). These connections could serve as a potential pathway for perceptual evidence accumulated in PPC to directly influence M1 excitability in situations in which motor plans are simple or are executed automatically (as in our experiment). This could explain the differences in the excitability of M1 that were observed in our experiment in trials with high PAS ratings, as more information would be transferred from PPC to M1.

### 4.4 Conclusions

Overall, our results shed new light on the relation between action and perceptual awareness by providing evidence that the motor system can be incorporated into metacognitive processes. Combined with previous results, these findings broaden our understanding of the interactions between action and conscious access that allow humans to dynamically adjust and re-evaluate their interactions with the environment. The significance of the influence of motor information on awareness judgments calls for broader theories of conscious access that primarily focus on processing sensory input.

## Author Contributions

Justyna Hobot, Michał Wierzchoń and Kristian Sandberg designed the study. Justyna Hobot programmed and performed the study. Justyna Hobot, Marcin Koculak and Borysław Paulewicz were involved in the data analysis. Justyna Hobot drafted the manuscript. Marcin Koculak provided changes to the manuscript. Michał Wierzchoń, Kristian Sandberg and Borysław Paulewicz provided comments on the manuscript. Justyna Hobot corrected the manuscript.

## Abbreviations

2AFC: two-alternative forced choice
PFC: prefrontal cortex
dlPFC: dorsolateral prefrontal cortex
FDI: first dorsal interosseous
M1: primary motor cortex
MEP: motor-evoked potential
MSO: maximal stimulator output
PAS: Perceptual Awareness Scale
PPC: posterior parietal cortex
PMd: dorsal premotor cortex
RMT: resting motor threshold
RTs: response times
S1: primary somatosensory cortex
SD: standard deviation
SEs: standard errors
TMS: transcranial magnetic stimulation.

## Acknowledgements

This article is based upon work from COST Action CA18106, supported by COST (European Cooperation in Science and Technology). We thank Nicolas McNair for helping to enable online TMS triggering by updating the MagPy Python package to work with the Magstim Super Rapid^2^ Plus^1^ stimulator. We thank Marta Siedlecka and Zuzanna Skóra for providing comments on our manuscript.

## Data Availability Statement

The raw data and the scripts for analyses are available at: https://osf.io/29n6j.

